# CircHIPK3 dysregulation of the miR-30c/DLL4 axis is essential for KSHV lytic replication

**DOI:** 10.1101/2021.10.07.463491

**Authors:** Katherine L. Harper, Timothy J. Mottram, Chinedu A. Anene, Becky Foster, Molly R. Patterson, Euan McDonnell, Andrew Macdonald, David Westhead, Adrian Whitehouse

## Abstract

Non-coding RNA (ncRNA) regulatory networks are emerging as critical regulators of gene expression. These intricate networks of ncRNA:ncRNA interactions modulate multiple cellular pathways and impact the development and progression of multiple diseases. Herpesviruses, including Kaposi’s sarcoma-associated herpesvirus, are adept at utilising ncRNAs, encoding their own as well as dysregulating host ncRNAs to modulate virus gene expression and the host response to infection. Research has mainly focused on unidirectional ncRNA-mediated regulation of target protein-coding transcripts; however, we have identified a novel host ncRNA regulatory network essential for KSHV lytic replication in B cells. KSHV-mediated upregulation of the host cell circRNA, circHIPK3, is a key component of this network, functioning as a competing endogenous RNA of miR-30c, leading to increased levels of the miR-30c target, DLL4. Dysregulation of this network highlights a novel mechanism of cell cycle control during KSHV lytic replication in B cells. Importantly, disruption at any point within this novel ncRNA regulatory network has a detrimental effect on KSHV lytic replication, highlighting the essential nature of this network and potential for therapeutic intervention.

## Introduction

Within the huge diversity of RNA species transcribed from the human genome, the majority have no coding capacity and are instead designated non-coding RNAs (ncRNAs). Even within ncRNAs, there is a wide repertoire of species generally categorised based on their size. Small ncRNAs, are less than 200 nucleotides in length, and include microRNAs (miRNAs), small-interfering, small nuclear, small nucleolar and PIWI-interacting RNAs [1]. In contrast, long ncRNAs (lncRNAs) vary in length from 200 nucleotides to 100 kilobases, and include long intergenic, long intronic, sense and antisense RNAs [2]. As interest in different ncRNAs species has increased, so too have their known assigned functions. It is now understood, that far being from errors in transcription, ncRNAs have key roles in regulating all aspects of cell biology and are critical regulators of gene expression. This is reinforced by ncRNA dysregulation being implicated in the development and progression of many disease states, including cancer, neurological disorders and infection [3] [4] [5].

To date research has mainly focused on unidirectional ncRNA-mediated regulation of target protein-coding transcripts, exemplified by miRNA-mediated degradation of mRNA [6]. However, there is emerging evidence of the existence of interplay between ncRNAs that strongly influence how gene expression is regulated by forming ncRNA regulatory networks. A cornerstone of these networks is the ability to act as competing endogenous RNA (ceRNAs), here ncRNAs can compete with each other for mRNA binding, thereby adding a new layer of regulatory potential. Several mechanistic interactions allow ncRNAs to function as ceRNA regulators including; miRNAs interacting with lncRNAs to reduce their stability, ncRNAs competing with miRNAs for the interaction with shared mRNA targets and ncRNAs acting as miRNA sponges or decoys to enhance target mRNAs expression [7] [8].

Circular RNAs (circRNAs) are a novel class of lncRNAs, characterised by a covalently closed loop lacking a 5’ end cap or 3’ Poly (A) tail. Originally thought to be rare splicing errors, recent breakthroughs have shown circRNAs are highly abundant and play key roles in gene regulation [9] [10]. circRNAs are mostly formed through a unique backsplicing mechanism, where a downstream splice donor is joined to an upstream splice acceptor forming a closed circle [11]. These structures are highly stable, conserved across many species and often cell-type or condition specific [12]. Implicated in a wide range of functions, including protein interactions and transcriptional regulators, interest in circRNAs was ignited with the emergence of their role as ceRNAs through miRNA interactions. In 2008, ciRS-7 was shown to sponge miR-7, acting as a key ncRNA regulator, since then numerous examples of functional ceRNA circRNAs have emerged, strongly implicated circRNAs as key regulators [13].

The ability of ncRNAs to act as regulators of gene expression, offers the potential for viruses to subvert these processes aiding in their own replication or to modulate the host cell response to infection. Herpesviruses, such as Kaposi’s Sarcoma-Associated Herpesvirus (KSHV), are particularly proficient at utilising ncRNAs regulatory pathways [14]. KSHV-encoded miRNAs and lncRNAs, as well as dysregulating host cell miRNAs to help in the establishment of a persistent latent infection, evading the immune response and regulating the switch between latent and lytic replication cycles [15] [16]. Recent evidence has also shown that KSHV encodes its own circRNAs, with a circ-vIRF4 identified by multiple studies [17] [18]. Hundreds of highly variable low copy number circRNAs are also expressed from the PAN locus, although functionality has not yet been established. Another circRNA derived from K12 gene contained the miRNA sequences also expressed from this gene, it has therefore been hypothesised it could act as ceRNA of its own miRNAs [19] This suggests viruses may dysregulate cellular circRNAs as part of complex ncRNA regulatory networks, aiding in viral manipulation of the host cell.

Herein we have identified that KSHV dysregulates a novel host cell ncRNA regulatory network which is essential for successful viral lytic replication and infectious virion production. circHIPK3 is a key component of this network and we have demonstrated that circHIPK3 acts as a ceRNA, sponging miR-30c leading to an increase in the levels of the miR-30c target, DLL4. We further demonstrate that disruption at any point within this novel network has a detrimental effect on KSHV lytic replication, highlighting the essential nature of this network.

## Results

### miR-29b and miR-30c are inhibitory miRNAs dysregulated during KSHV lytic replication

To determine whether KSHV infection dysregulates specific host cell ncRNA networks, we first assessed which cellular miRNAs were dysregulated during KSHV reactivation in B cells, as miRNAs are central in ncRNA axes. miR-Seq [20] was utilised to identified miRNAs which are altered during the course of KSHV lytic replication at 0, 16 and 24 hours post reactivation in TREx-BCBL1-RTA cells, a KSHV-latently infected B-lymphocyte cell line containing a Myc-tagged version of the viral RTA under the control of a doxycycline-inducible promoter. Z-score analysis of the miR-Seq highlighted the most consistently dysregulated miRNAs, with the vast majority downregulated during lytic replication in B cells (Fig. 1a). Based on a combination of known function, abundance and Z-score, miR-30c and miR-29b were selected for further investigation. This was further supported by available microarray online datasets showing miR-29b and miR-30c are downregulated in KS patient clinical samples compared to levels in adjacent tissues (Fig. 1b and c) and in KSHV positive B cell lines (Fig. S1) [21]. In addition, the miR-30 family has been previously observed to be downregulated upon KSHV infection of lymphatic endothelial cells [22]. Other miRNAs highlighted in our miR-Seq were not significantly changed in online microarray datasets, further prioritizing selection of miR-29b and miR-30c for further investigation (Fig. S2). To conclusively confirm downregulation of miR-29b and miR-30c in TREx-BCBL1-RTA cells, qPCR analysis was utilised to determine mature miRNA levels. Results confirmed a dramatic reduction of mature miR-29b and miR-30c levels at 24 hours post reactivation, at 75% and 70% respectively (Fig. 1d). Notably, quantification of the primary and pre-miRNAs levels via qPCR showed no significant decrease, implicating downregulation occurs at the miRNA mature levels, potentially implicating ncRNA network regulation (Fig. 1e and f).

**Figure 1:**
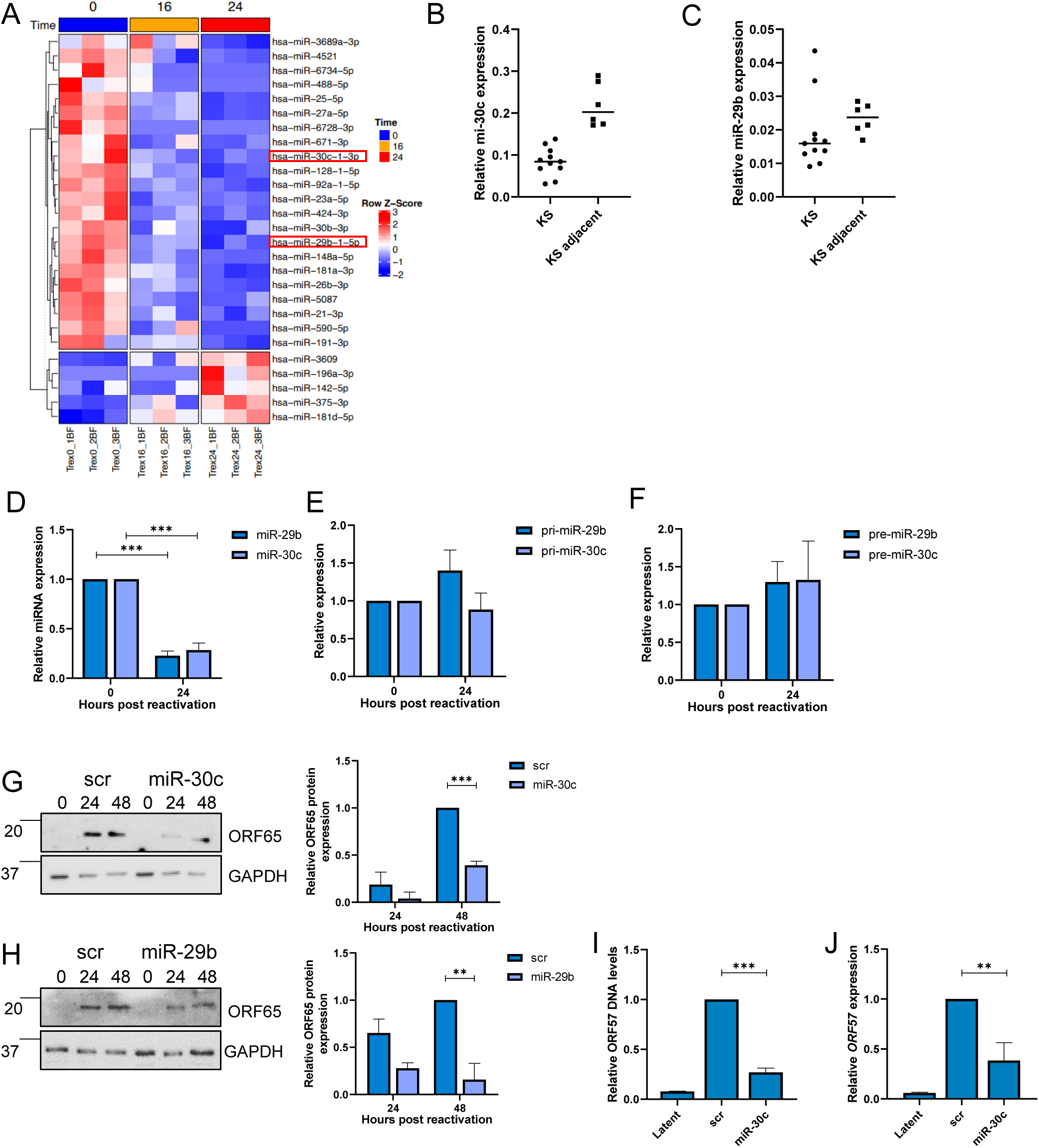
KSHV dysregulates inhibitory miRNA. **(A)** Heat map of differentially expressed miRNAs during KSHV replication at 0, 16 and 24 hour identified through miR-Seq. miR-29b and miR-30c are highlighted in red, with T test analysis. **(B)** Scatter plot data of miR-30c from GSE55265 with KS samples (n=11) and KS-adjacent samples (n=6). **(C)** Scatter plot data of miR-29b from GSE55265 with KS samples (n=11) and KS-adjacent samples (n=6). **(D)** qPCR analysis of miR-29b and miR-30c levels in cells at 0 and 24 hours post-lytic induction. SNORD68 was used as a housekeeper (n=3). qPCR analysis of TREx 0 and 24 post-induction (n=3) using GAPDH as a housekeeper, analysis is of levels of pri-miR-29b/pri-miR-30c **(E)** and pre-miR-29b/pre-miR-30c **(F). (G)** Representative western blot of scr or miR-30c transfected TREx cells analysed for ORF65 expression with GAPDH as a loading control. Densitometry analysis performed on n=3. **(H)** Representative western blot of scr or miR-29b transfected TREx cells analysed for ORF65 expression with GAPDH as a loading control. Densitometry analysis performed on n=3. **(I)** qPCR analysis of ORF57 DNA levels for viral load at 72 hours post induction, with scr and miR-30c transfected cells with uninduced cells as a control and GAPDH as a housekeeper (n=3). **(J)** qPCR analysis of *ORF57* in HEK-293T cells for reinfection assay. Error bars represent standard deviation, *P<0.05, **P<0.01, ***P<0.001 (Student’s t-test).

Due to the consistent downregulation we speculated that miR-29b and miR-30c may have an inhibitory effect on KSHV lytic replication in B cells. miRNA mimics were therefore transfected separately into TREx-BCBL1-RTA cells resulting in overexpression of miR-29b and miR-30c (Fig. S3). Reactivation assays demonstrated that miR-29b and miR-30c overexpression resulted in a significant reduction in the lytically expressed late ORF65 protein, compared to a scrambled control (Fig. 1g and h). Further experiments focused on miR-30c due to its higher expression levels. Additionally, viral genomic DNA was measured via qPCR from scrambled and miR-30c overexpressing TREx-BCBL1-RTA cells to assess whether viral DNA load was affected, with cells overexpressing the miRNA having a 75% reduction (Fig. 1i). To examine whether miR-30c overexpression also affected infectious virion production, supernatants of reactivated scrambled and miRNA overexpressing TREx-BCBL1-RTA cells were used to re-infect naïve HEK-293T cells and qPCR used to determine KSHV ORF57 expression. Cells reinfected with supernatant from miRNA overexpressing cells contained 60% less viral RNA compared to controls (Fig. 1j). Taken together these data suggest that KSHV lytic replication in B cells is impacted by the overexpression of miR-29b and miR-30c, suggesting they are downregulated during KSHV lytic replication due to their inhibitory effect.

### circHIPK3 sponges miR-29b and miR-30c

We next determined whether KSHV could dysregulate these inhibitory miRNAs through a ncRNA network. CircRNAs, have previously been reported to act as miRNA sponges thereby regulating miRNA/mRNA axes. Notably, RNA binding predictions highlighted circHIPK3 (hsa_circ_0000284) contained complementary sequences to both miR-29b and miR-30c, suggesting it may function as a potential sponge for both miRNAs (Fig. S4). To assess whether circHIPK3 levels were altered during KSHV lytic replication, qPCR was performed using divergent primers annealing at the distal ends of circHIPK3. Results showed a ~ 3-fold increase of circHIPK3 at 24 hours post reactivation compared to latent samples (Fig. 2a). Notably, this upregulation was specific to the circular transcript, as the mRNA levels of *HIPK3* showed no significant change (Fig. 2b). Given the potential of circHIPK3 to function as a miR-29b and miR-30c sponge during KSHV lytic replication, we next assessed its sub-cellular localisation. Fluorescent *In Situ* hybridisation (FISH), using specific probes against the unique backsplice site within circHIPK3, demonstrated a clear cytoplasmic localisation in TREx-BCBL1-RTA cells (Fig. 2c). To further support a potential sponging activity of circHIPK3, Ago2 RIPs were performed to determine whether circHIPK3 associates with the miRNA machinery. Results show a significant enrichment of circHIPK3 in the Ago2 RIP over the IgG control (Fig. 2d). Moreover, overexpression of either miR-29b or miR-30c mimics further enhanced this enrichment in the Ago2 RIP (Fig. 2d). Finally, to confirm direct binding between circHIPK3 and miR-29b or miR-30c, biotinylated miR-29b or miR-30c were transfected into HEK-293Ts and a RIP was performed using Streptavidin beads. Results showed a significant enrichment of circHIPK3 in the miRNA transfected samples compared to scrambled controls (Fig. 2e). Together these data suggests that circHIPK3 is upregulated during KSHV lytic replication and functions as a molecular sponge for miR-29b and miR-30c.

**Figure 2:**
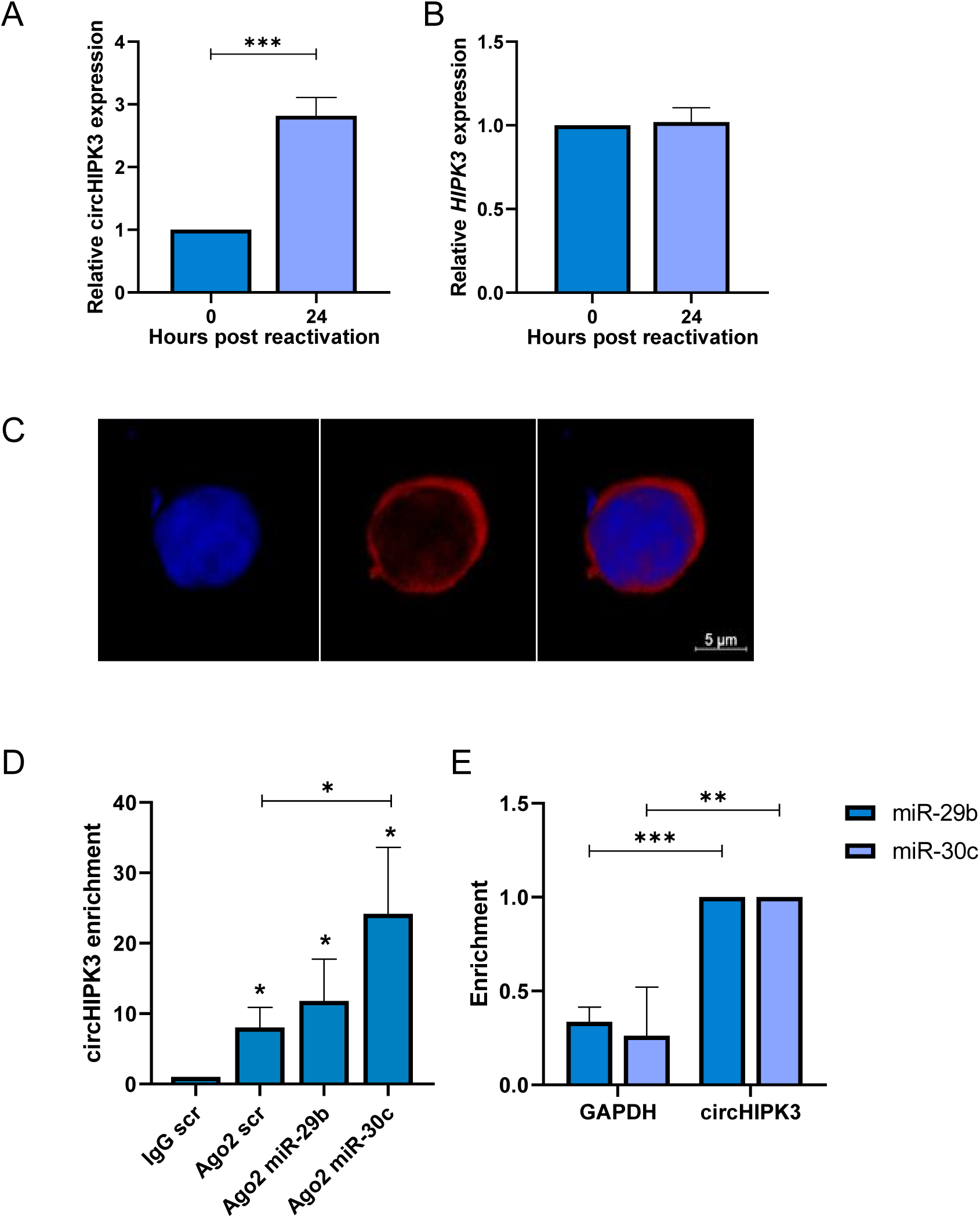
circHIPK3 sponges miR-29b and miR-30c. qPCR analysis of TREx 0 and 24 post-induction (n=3) using GAPDH as a housekeeper, analysis is of levels of circHIPK3 **(A)** and *HIPK3* **(B). (C)** FISH analysis of TREx cells with probes against circHIPK3 (red). DAPI was used as nuclei stain (blue), scale bar represents 5 μM. **(D)**. qPCR analysis of Ago2 RIPs in TREx cells transfected with scr, miR-29b or miR-30c showing circHIPK3 enrichment over GAPDH (n=3). **(E)** qPCR analysis of circHIPK3 or GAPDH enrichment in HEK-293T cells transfected with either biotinylated miR-29b or miR-30c over scr (n=3). Error bars represent standard deviation, *P<0.05, **P<0.01, ***P<0.001 (Student’s t-test).

### CircHIPK3 is essential for KSHV replication

To assess the importance of circHIPK3 on KSHV lytic replication, TREx-BCBL1-RTA cells were transduced with lentivirus-based shRNAs targeting the unique backsplice of circHIPK3. qPCR confirmed successful depletion of circHIPK3, but importantly knockdown had little effect on the *HIPK3* transcript (Fig. 3a and b), meaning any effects of the knockdown was solely due to circHIPK3 depletion. Reactivation assays demonstrated that circHIPK3 depletion resulted in a significant reduction in ORF65 protein production compared to the scrambled control (Fig. 3c). The effect of circHIPK3 knockdown was further evaluated using viral DNA load and reinfection assays, with a significant decrease in both viral load and infectious virion production being observed upon depletion (Fig. 3d and e). Together this confirms that KSHV specifically requires the function of circHIPK3 to undergo efficient lytic replication and infectious virion production. With the importance of circHIPK3 of KSHV lytic replication confirmed, we next assessed what effect circHIPK3 depletion had upon levels of miR-29b or miR-30c during lytic replication. As previously observed both miR-29b and miR-30c were downregulated in scrambled control cells at 24 hours post reactivation, in contrast no significant downregulation was observed in miR-29b and miR-30c levels upon circHIPK3 depletion (Fig. 3f). This effect was specific to miR-29b and miR-30c as circHIPK3 depletion had no effect on miR-27a levels, another miRNA downregulated upon KSHV lytic replication (Fig. 3f). Together this shows that KSHV-mediated circHIPK3 dysregulation can specifically affect miR-29b and miR-30c levels.

**Figure 3:**
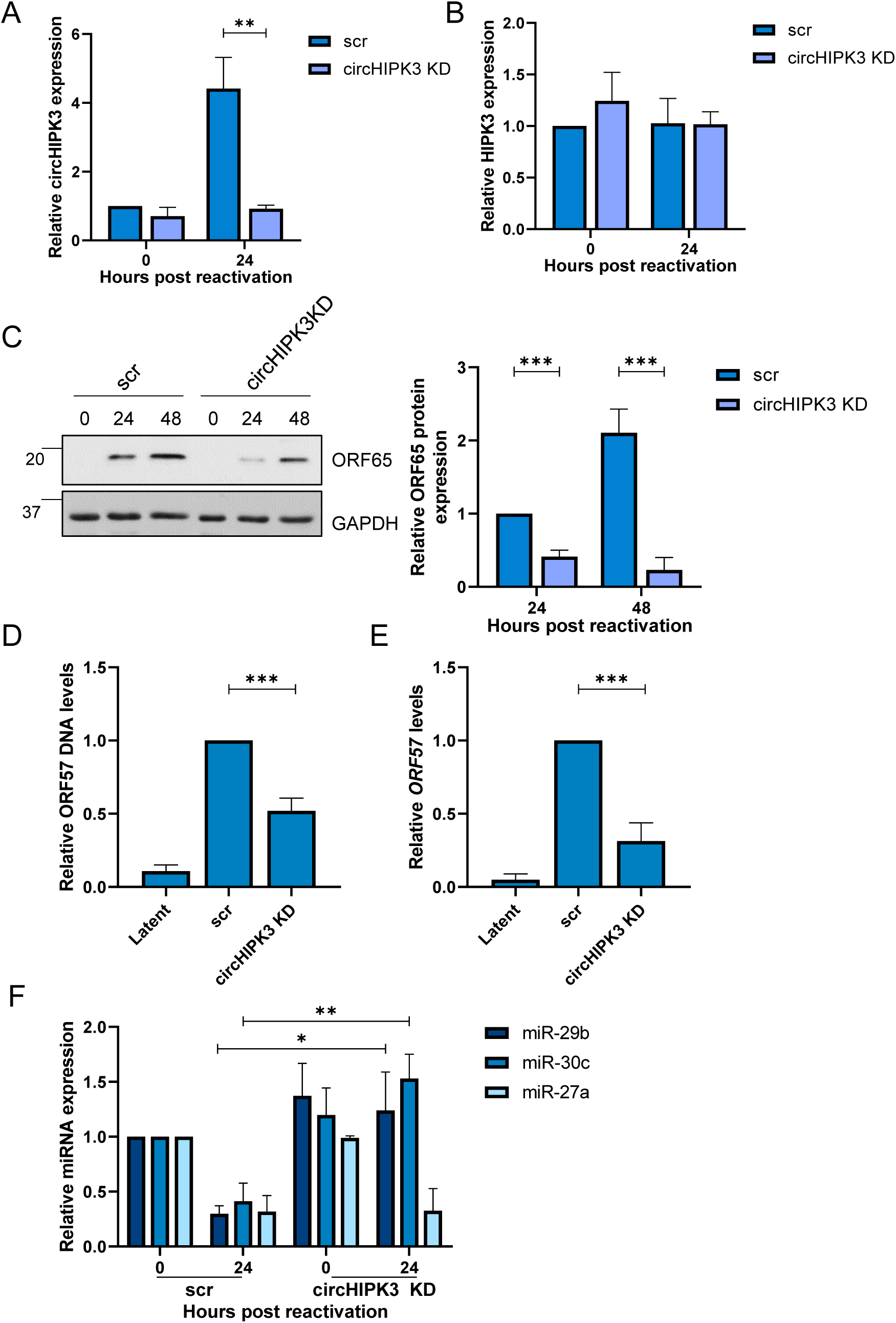
circHIPK3 is essential for KSHV replication. **(A)** qPCR analysis of circHIPK3 levels in scr and circHIPK3 KD stably expressing TREx cells. GAPDH was used as a housekeeper (n=3). **(B)** qPCR analysis of *HIPK3* levels in scr and circHIPK3 KD stably expressing TREx cells. GAPDH was used as a housekeeper (n=3). **(C)** Representative western blot of ORF65 levels in scr and circHIPK3 KD cell lines with GAPDH as a loading control. Densitometry analysis of n=3 western blots. **(D)** qPCR analysis of ORF57 DNA levels for viral load at 72 hours post induction, with scr and circHIPK3 KD cells with uninduced cells as a control and GAPDH as a housekeeper (n=3). **(E)** qPCR analysis of *ORF57* in HEK-293T cells for reinfection assay. **(F)** qPCR analysis of levels of miR-29b, miR-30c and miR-27a in scr and circHIPK3 KD cells 0 and 24 hours post-induction. SNORD68 was used as a housekeeper (n=3). Error bars represent standard deviation, *P<0.05, **P<0.01, ***P<0.001 (Student’s t-test)

### KSHV ORF57 expression enhances circHIPK3 levels

To determine if any KSHV-encoded proteins are involved in dysregulating circHIPK3 levels, we firstly assessed circHIPK3 levels at different time points post reactivation. qPCR showed a clear upregulation between 16 and 24 hours (Fig. 4a). We therefore examined whether any early KSHV proteins were sufficient to induce circHIPK3 levels independently. HEK-293T cells were transfected with control GFP, or KSHV ORF50-GFP and ORF57-GFP expression constructs [23] [24] and circHIPK3 levels assessed by qPCR at 24 hours post transfection. Results show expression of ORF57-GFP alone was sufficient to increase circHIPK3 levels (Fig. 4b) and this upregulation was dose-dependent (Fig. 4c). Western blotting confirmed successful dose-dependent transfections (Fig. S5a and S5b). KSHV ORF57 is a multifunctional RNA binding protein involved in several stages of viral RNA processing [25] [26]. Therefore, we assessed whether ORF57 interacted with circHIPK3 using RNA immunoprecipitations performed in control or ORF57-GFP transfected HEK-293T cells utilising GFP-TRAP beads. RIPs were performed at 16 and 24 hours post-transfection and results showed that ORF57 associated with both circHIPK3 and the linear HIPK3 transcript strongly at 16 hours. In contrast, at 24 hours ORF57 showed stronger association with circHIPK3 compared to the linear form (Fig. 4d). This suggests that ORF57 binds both the linear and circular forms of HIPK3 and may have a role in the promotion of circularisation of linear HIPK3 transcript. (Fig. 4d). The role of ORF57 was further investigated with previously characterised RNA binding and dimerisation negative mutants [23] [24], with the RNA binding mutant unable to bind to circHIPK3 (Fig. S6), however, the dimerisation mutant was too unstable to utilise (Fig. S7). Together these data suggest that ORF57 may be involved in the increase in circHIPK3 levels during KSHV lytic replication.

**Figure 4:**
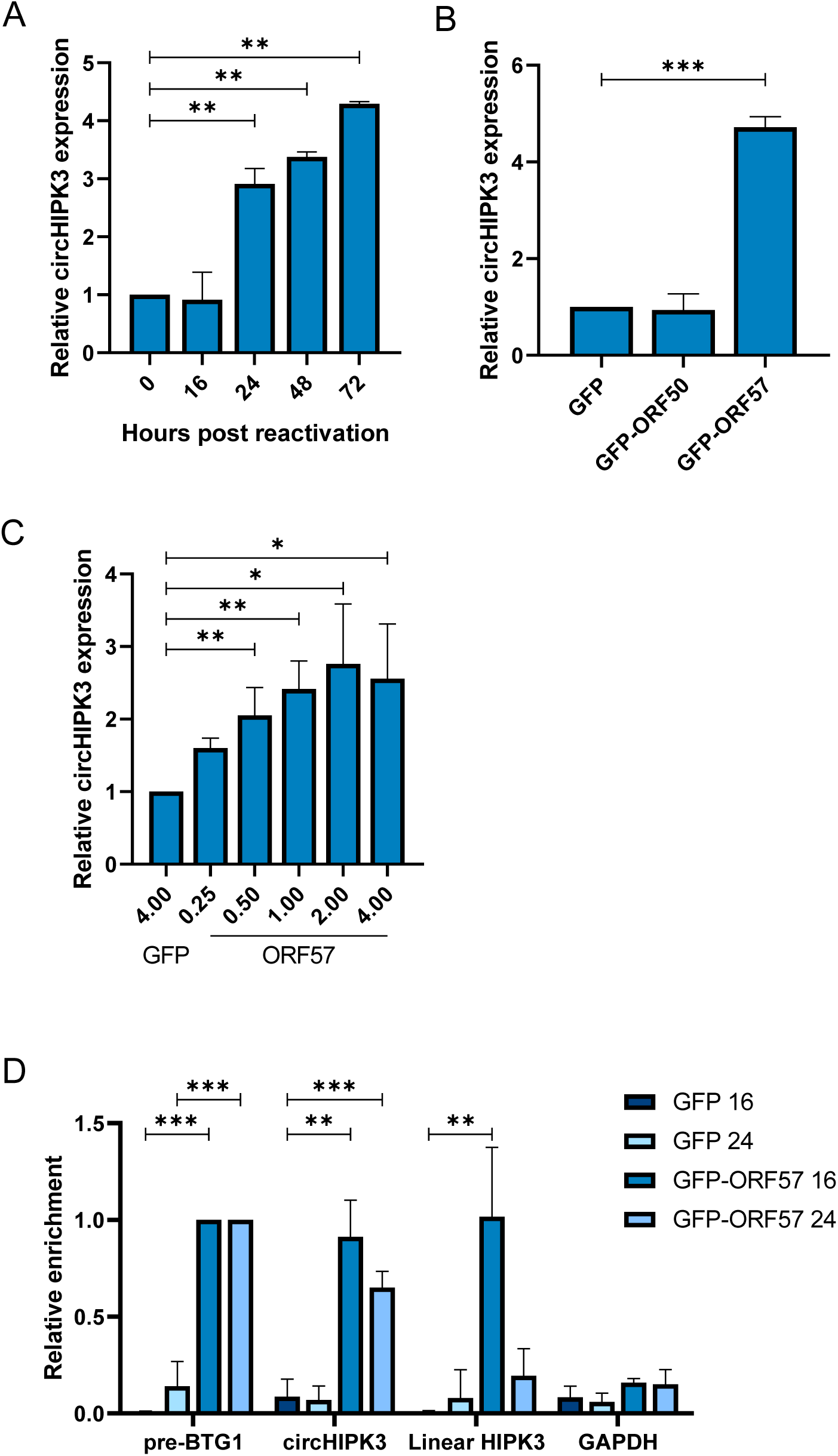
ORF57 contributes to the upregulation of circHIPK3 levels. **(A)** qPCR analysis of circHIPK3 levels at time points in TREx post-induction with GAPDH as a housekeeper (n=3). **(B)** qPCR analysis of circHIPK3 expression in HEK-293Ts transfected with GFP, ORF50-GFP or ORF57-GFP with GAPDH as a housekeeper (n=3). **(C)** qPCR analysis of circHIPK3 expression in HEK-293Ts transfected with 4 µg GFP, or 0.25-4 µg ORF57-GFP, GAPDH was used as a housekeeper (n=3). **(D)** qPCR analysis GFP RIPs in GFP or ORF57-GFP transfected HEK-293Ts at 16 and 24 hours post-transfection respectively n=3 for each time course. Error bars represent standard deviation, *P<0.05, **P<0.01, ***P<0.001 (Student’s t-test)

### DLL4 is regulated by miR-30c and circHIPK3

The dependence of the circHIPK3/miR-30c axis to enhance KSHV lytic replication in B cells, suggests that KSHV may modulate this ncRNA axis to regulate expression of miR-30c-specific mRNA targets. Previous results has shown that the miR-30 family targets DLL4 [22] and combined cross referencing of online databases with upregulated mRNAs from RNA Seq during KSHV infection (Fig. S8), qPCR and immunoblotting confirmed DLL4 was upregulated at both the RNA and protein level during KSHV lytic replication in TREx-BCBL1-RTA cells (Fig. 5a and 5b). To determine whether the circHIPK3/miR-30c axis specifically affected DLL4 expression, we assessed what effect either circHIPK3 depletion or overexpression of a miR-30c mimic had on DLL4 expression in TREx-BCBL1-RTA cells. In both cases we observed a significant reduction in DLL4 mRNA and protein levels, implying both circHIPK3 and miR-30c regulate DLL4 levels (Fig. 5c-f). These results were further supported using a 3’ UTR luciferase assay, where the 3’ UTR of DLL4 was cloned into a luciferase reporter plasmid. Transfection of miR-30c lead to a reduction in luminescence, which confirmed miR-30c directly regulates DLL4 expression in TREx-BCBL1-RTA cells (Fig. 5g).

**Figure 5:**
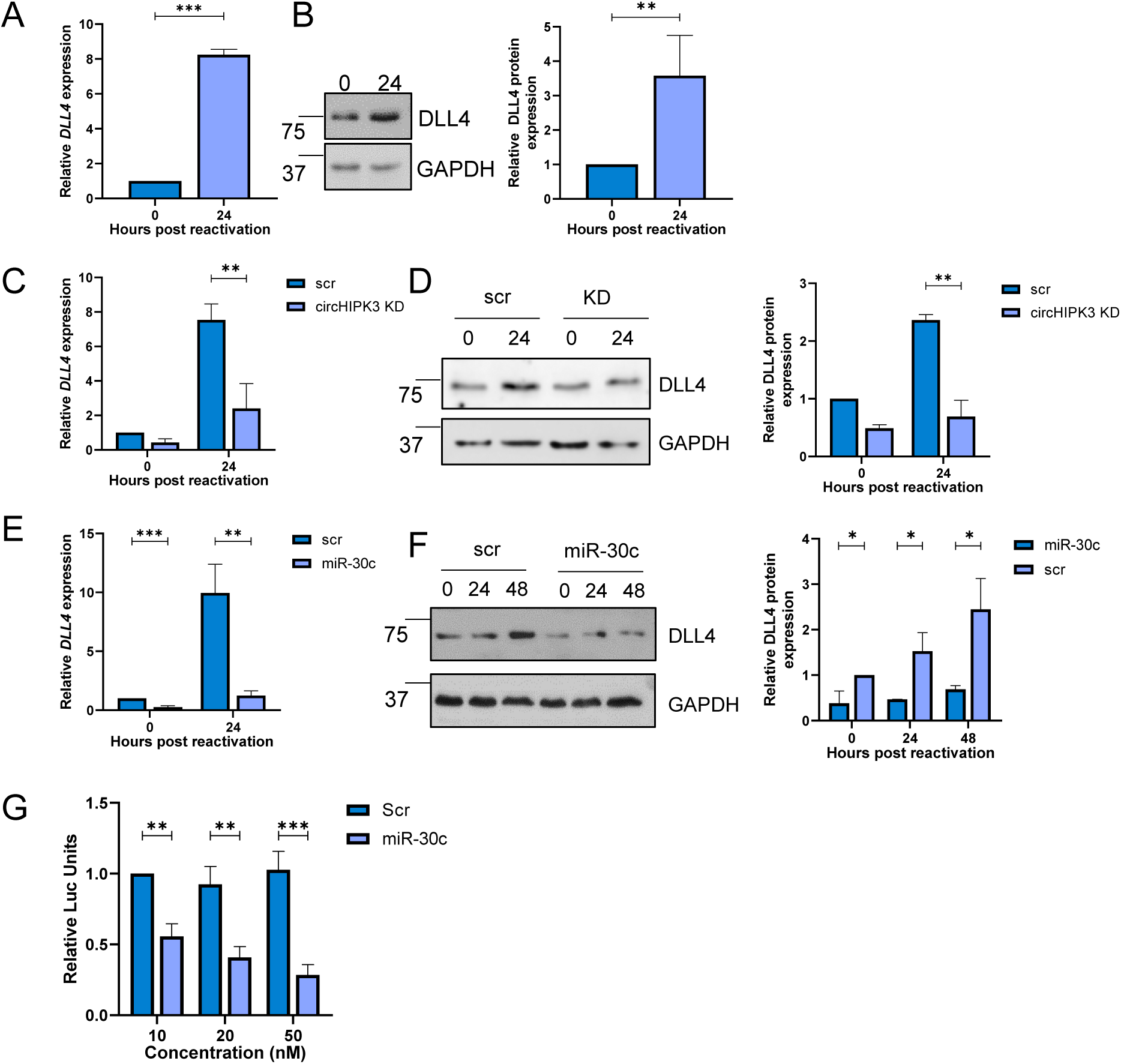
DLL4 is targeted by circHIPK3 and miR-30c. **(A)** qPCR analysis of *DLL4* levels in TREx cells 0 and 24 hours post induction. GAPDH was used a housekeeper (n=3). **(B)** Representative western blot of DLL4 expression in TREx cells 0 and 24 hours post induction. GAPDH was used a loading control (n=3). **(C)** qPCR of *DLL4* levels in scr and circHIPK3 KD stable TREx cells at 0 and 24 hours post induction with GAPDH as a housekeeper (n=3). **(D)** Representative western blot of DLL4 levels in scr and circHIPK3 stable TREx cells at 0 and 24 hours post induction with GAPDH as a loading control. Densitometry analysis of n=3. **(E)** qPCR analysis of *DLL4* levels in TREx cells that have been transfected with either scr of miR-30c 24 hours pre-induction, analysis performed 0 and 24 hours post induction. GAPDH was used a housekeeper (n=3). **(F)** Representative western blot of DLL4 levels in scr or miR-30c transfected TREx cells at 0 and 24 hours post induction with GAPDH as a loading control. Densitometry analysis of n=3 **(G)** Luciferase Reporter Assay from HEKL-293Ts co-transfected with DLL4 3’UTR reporter plasmid and either scr or miR-30c. Data presented is relative to an internal firefly control (n=4). *P<0.05, **P<0.01, ***P<0.001 (Student’s t-test)

### DLL4 is essential for KSHV lytic replication in B cells

To confirm the importance of DLL4 during KSHV lytic replication, TREx-BCBL1-RTA cells were transduced with lentivirus-based shRNAs, significantly depleting DLL4 RNA and protein expression during lytic replication (Fig. 6a and 6b). Reactivation assays demonstrated that DLL4 depletion resulted in a significant reduction in late ORF65 protein expression compared to the scrambled control (Fig. 6c). Similarly, DLL4 depletion resulted in a significant reduction in viral load and infectious virion production, compared to scrambled control (Fig. 6d and e). Together, these data highlight that dysregulation of a circHIPK3/miR-30c/DLL4 regulatory circuit is essential for efficient KSHV lytic replication.

**Figure 6:**
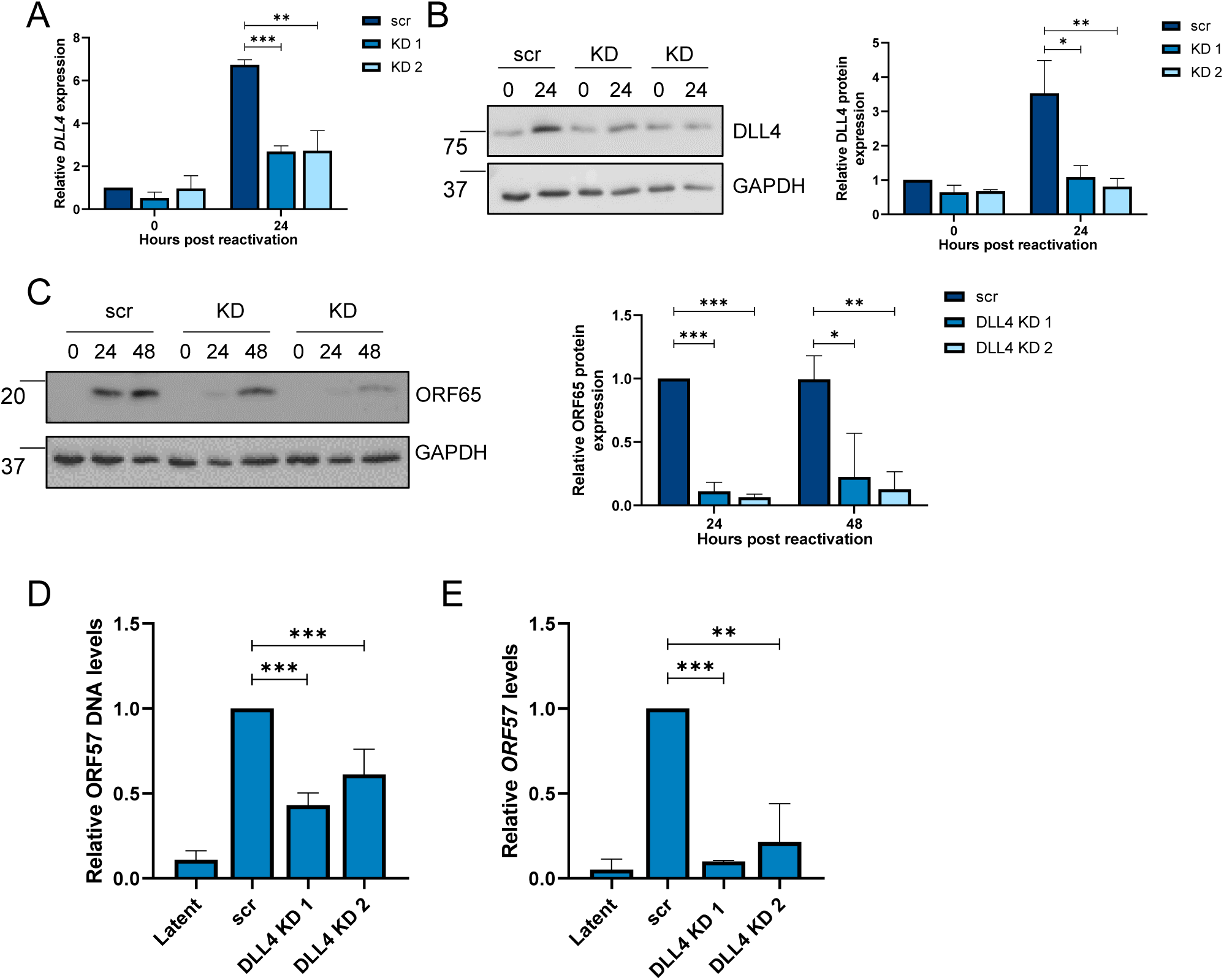
circHIPK3:miR-30c:DLL4 dysregulation affects cell cycle. **(A**) qPCR of *DLL4* expression in stable expression scr and 2 *DLL4* KD TREx cell lines at 0 and 24 hours post-induction. GAPDH was used as a housekeeper (n=3). **(B)** Representative western blot of DLL4 levels in scr and DLL4 KD TREx cells with GAPDH as a loading control, densitometry analysis is n=3. **(C)** Representative western blot of ORF65 levels at 0, 24 and 48 hours post induction in scr and DLL4 KD TREx cells, GAPDH was used as a loading control and densitometry analysis performed on n=3. **(D)** qPCR analysis of ORF57 DNA levels for viral load at 72 hours post induction, with scr and DLL4 KD cells with uninduced cells as a control and GAPDH as a housekeeper (n=3). **(E)** qPCR analysis of *ORF57* in HEK-293T cells for reinfection assay. *P<0.05, **P<0.01, ***P<0.001 (Student’s t-test)

DLL4 is a transmembrane protein that acts as a ligand for Notch receptors 1 and 4. As such, it has multiple functions, for instance modulating endothelial cell behaviour during angiogenesis [22] and regulating cell cycle-associated proteins [27]. Focusing on cell cycle-associated proteins, we analysed several cyclins during KSHV lytic replication in TREx-BCBL1-RTA cells, comparing scrambled control, circHIPK3 KD and DLL4 KD cell lines. Lytic replication led to dysregulation of both *CCNE1* and *CCNB1*, whereas this downregulation was partly abrogated in both circHIPK3 and DLL4 KD cell lines (Fig. 7a and b). As KSHV DNA replication is tied to viral manipulation of the cell cycle and our previously observed reductions in viral loads in both circHIPK3 and DLL4 KD cell lines, we assessed what effect dysregulation of the circHIPK3/miR-30c/DLL4 regulatory circuit had upon KSHV-mediated cell cycle dysregulation during lytic replication. Flow cytometry analysis using propidium iodide demonstrated that scrambled control TREx-BCBL1-RTA cells showed a significant drop in the number of cells in G2/M phase during lytic replication and a maintained number in G1 phase. In contrast, depletion of either circHIPK3 or DLL4 resulted in an increased number of cells in G2/M phase (Fig. 7c and d). Together this data suggests that dysregulation of the circHIPK3/miR-30c/DLL4 regulatory circuit affects cell cycle regulation to enhance KSHV lytic replication. This is clearly supported by results showing disruption of this axis at several points can inhibit virus replication. To further confirm KSHV-mediated dysregulation of the cell cycle is important for viral DNA replication, cells were treated with cell-cycle inhibitors RO-3306, nocodazole and thymidine and effects on the cell cycle were confirmed with flow cytometry (Fig. S9). Treatment with the G2/M phase arresting RO-3306 and nocodazole, led to a significant decrease in viral load in KSHV lytic cells (Fig. 7e). In contrast, treatment with thymidine, a G1/S exit blocker, led to no decrease in viral load in scrambled control cells. However, more importantly thymidine treatment of DLL4 KD cells, reversing DLL4-mediated cell cycle changes led to rescue of viral DNA replication (Fig. 7f). This suggests that the effect of DLL4 on KSHV lytic replication in B cells are directly mediated through its cell cycle activities. Cell cycle dysregulation has been shown to be important for KSHV DNA replication, and we confirmed this with both the knockdown cell lines and cell cycle blocking inhibitors. Therefore, we hypothesised that due to this inhibition of viral DNA replication, formation of viral replication centres, where viral DNA replication occurs, would be reduced or delayed. To investigate this further, confocal microscopy was performed to examine whether formation of viral replication centres were disrupted in circHIPK3 and DLL4 KD cell lines, with a significant reduction observed (Fig. 7g). Together this shows viral DNA replication is impaired in both circHIPK3 and DLL4 KD cell lines and through treatment with cell cycle inhibitors. Together these data suggest KSHV-mediated dysregulation of the circHIPK3/miR30c/DLL4 axis is a novel mechanism for cell cycle regulation in B cells.

**Figure 7:**
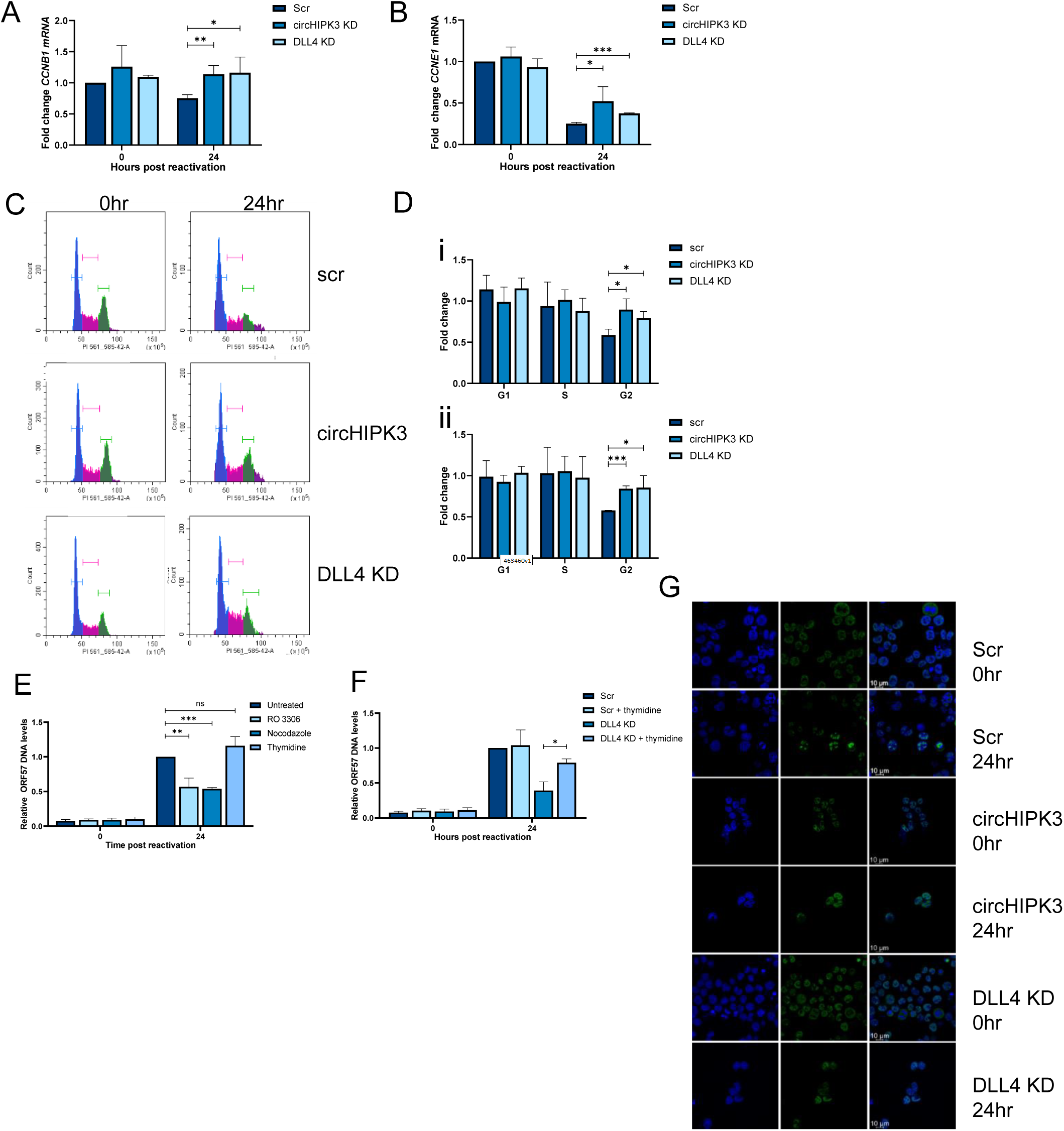
circHIPK3:miR-30c:DLL4 dysregulation affects cell cycle. **(A)** and **(B)** qPCR of *CCNB1* and *CCNE1* levels in scr, circHIPK3 KD and DLL4 KD cells at 0 and 24 hours post induction (n=3). **(C)** Representative plot of cell cycle distribution at 48 hours post induction in scr, circHIPK3 and DLL4 KD cells, with G1 (blue), S (pink) and G2 (green) highlighted. **(D)** Analysis of fold change in number of cells in G1, S and G2 phases of the cell cycle from 0 to 24 hours post-induction (i) or 0 to 24 hours post-induction (ii). Cells used were scr, circHIPK3 KD or DLL4 KD stable TREx cells and cycle analysis was performed using propidium iodide and flow cytometry (n=3). **(E)** qPCR for ORF57 DNA levels at 48 hours post lytic induction with treatment of RO 3306, nocodazole or thymidine. **(F)** qPCR for ORF57 DNA levels at 48 hours post lytic induction with treatment of RO 3306, nocodazole or thymidine in scr or DLL4 KD cell lines. **(G)** Representative immunofluorescence of scr, circHIPK3 KD or DLL4 KD cell lines at 0 and 24 hours. Sections are stained for DAPI (blue) and RNA pol II (green). Replication centres are highlighted with white arrow. Scale bars represent 10 µ M. *P<0.05, **P<0.01, ***P<0.001 (Student’s t-test)

**Figure 8:**
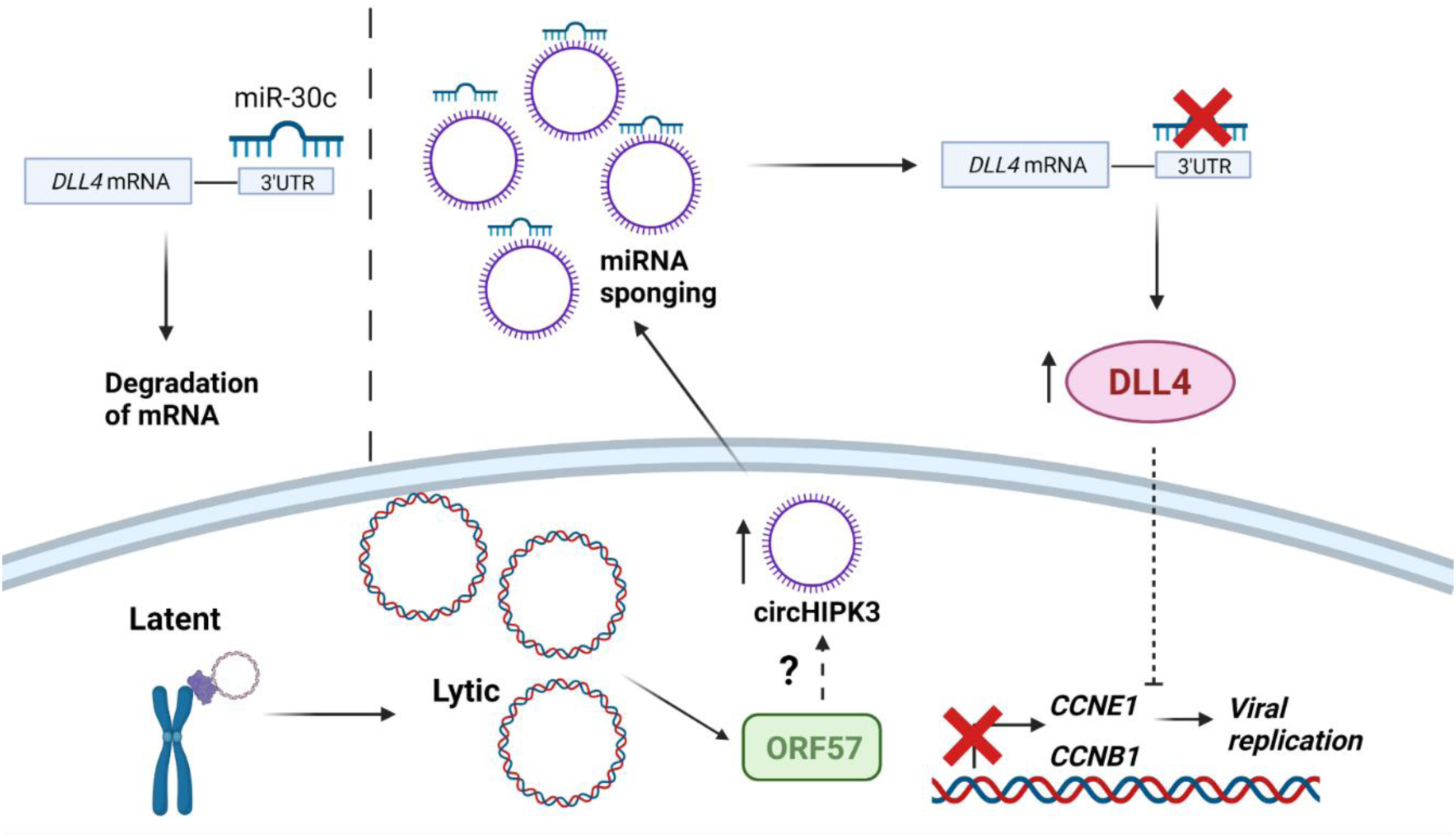
KSHV dysregulates a novel ncRNA network during lytic replication. circHIPK3 is upregulated by KSHV, partly mediated through ORF57, this leads to increased sponging of miR-30c and increased DLL4 levels. DLL4 has downstream affects on cyclins leading to changes in the cell cycle aiding KSHV replication.

## Discussion

Despite circRNAs relatively recent discovery, functional analysis has suggested a wide range of crucial roles in cell regulation including transcriptional regulation, miRNA sponging and protein regulation [28] [29]. It is not surprising therefore the dysregulation of circRNAs is implicated in a range of diseases including diabetes, dementia and cancer [30] [31] [32]. Here we show that the dysregulation of circRNAs also play a role in virus infection. We demonstrate that circHIPK3 is consistently upregulated during KSHV lytic replication and crucially, reversion of this upregulation leads to inhibition of viral replication and infectious virion production, highlighting the importance to KSHV replication. Furthermore, we performed functional analysis of the role circHIPK3 has during viral replication and have identified it as the first step in a novel non-coding RNA regulatory network that is dysregulated during KSHV infection.

Previous research has identified circHIPK3 is capable of miRNA sponging, including miR-7, miR-124 and another member of the miR-30 family: miR-30a [33] [34] [35]. Many of these miRNAs are classified as tumour suppressive, implicating circHIPK3 as a pro-proliferative factor, moreover, circHIPK3 is found upregulated in several cancers, including bladder, lung and HCC, and is therefore hypothesised to have regulatory roles in cell proliferation, cell cycle regulation and invasion [36] [37] [38]. However, the role circHIPK3 has not previously been elucidated in viral infection. KSHV has been shown to be prolific at utilising ncRNAs to aids its replication and propagation, with its complex genome encoding both miRNAs and circRNAs [17] [39], while dysregulation of cellular miRNAs enables low-immunogenicity post-transcriptional regulation of many different targets. Through dysregulation of circRNAs, such as circHIPK3, it enables upstream regulation of these miRNAs and through this ncRNA axis, allows subversion of cellular processes aiding viral replication and tumourigenesis. Similarly, a circARFGEF1/miR-125a-3p/GLRX3 axis has been shown to be required for KSHV vIRF1 induction of cell motility, proliferation and *in vivo* angiogenesis [40].

CircRNA functionality is tied to their localisation, with nuclear retained circRNAs interacting with RBPs or acting as transcriptional regulators [41] [42], whereas using RNA FISH, circHIPK3 was shown to be localised in the cytoplasm, suggestive of a role in miRNA sponging. This was further supported by RIPs of biotinylated miR-29b and miR-30c. These RIPs showed enrichment of circHIPK3, confirming that circHIPK3 acts as a competitive endogenous RNA and binds to these miRNAs. circHIPK3 was also enriched in Ago2 RIPs, implying it is interacting with the miRNA machinery. This once again supports its sponging function, with other famous miRNA-sponging circRNAs such as circSRY and circCDR1 also associating with Ago2 [43]. Of particular note is this enrichment was enhanced after transfection of miR-29b and miR-30c, suggesting when there are increased levels of target miRNAs interacting with Ago2, there is increased interaction by the sponging circRNA. Possibly either as the sponging circRNA is also upregulated as a compensatory mechanism or increased levels of the target miRNA can increase the stability of the interaction with Ago2, acting as a bridge.

Overexpression of miR-29b and miR-30c mimics had an inhibitory effect on KSHV lytic replication, therefore we hypothesise KSHV upregulates circHIPK3 to reduce the levels of these miRNAs. Crucially circHIPK3 depletion both inhibited viral replication, viral load and infectious virion production and also led to increases in miR-29b and miR-30c levels, implying a regulatory axis. Notably, analysis confirmed miR-29b and miR-30c downregulation did not occur at the primary or pre stages of miRNA biogenesis, once again implicating circHIPK3 as a miRNA sponge. miR-30c was selected for further investigation over miR-29b, due to its higher expression levels and increased circHIPK3 association with Ago2 in the presence of miR-30c. circHIPK3 depletion or overexpression of the miRNAs led to inhibition of KSHV lytic replication clearly demonstrating the importance of cellular circRNA dysregulation to enable virus-mediated hijacking of cellular ncRNA networks enhancing viral replication.

The mechanism behind this dysregulation is not fully elucidated. However, our analysis suggests that the KSHV lytic protein, ORF57, plays a significant role. Initial investigations show circHIPK3 dysregulation occurs between 16 and 24 hours upon lytic replication, coinciding with ORF57 expression during the viral cascade. Furthermore, ORF57 expression alone increased circHIPK3 levels. Interestingly, RIPs performed at 16 and 24 hours after transfection indicated ORF57 bound to *HIPK3* and circHIPK3 with the ratios changing over time, suggesting a direct role in regulating the splicing. At present, the actual mechanism of ORF57 mediated upregulation requires further investigation with cellular factors still to be identified. ORF57 is a promiscuous RBP, with many interacting partners, as transfection of ORF57 into HEK-293T cells lead to an increase in circHIPK3 levels without any other viral proteins, it is likely ORF57 recruits and utilises cellular proteins to aid this dysregulation, however, as of yet these factors are unknown. Furthermore, we show its RNA binding properties are likely important in this role, with the ORF57 RNA binding negative mutant RGG1/2 unable to bind to circHIPK3 (Fig. S6). Interestingly ORF57 is known to form a dimer [44]; many of the other known circRNA biogenesis promoting proteins act through dimerisation, for instance QKI and Mbl [45] [46]. Here the protein monomers bind to the upstream and downstream splice sites, and as they dimerise they bring the splice sites in close contact promoting circularisation, it is possible that ORF57 promotes circHIPK3 biogenesis in this form. Unfortunately the dimerisation negative ORF57 mutant we utilised was too unstable for further investigation (Fig. S7) [47].

DLL4 was identified as a potential target of the circHIPK3/miR-30c ncRNA regulatory axis. Supporting this potential network, we have shown that DLL4 is upregulated following KSHV lytic reactivation. Importantly, upon circHIPK3 depletion and miR-30c overexpression, we show, DLL4 upregulation is significantly reduced. Moreover, DLL4 depletion has an inhibitory effect on viral replication, affecting late ORF65 protein production, viral load and infection virion production. The importance of DLL4 to KSHV replication has been previously investigated, with the viral protein vGPCR upregulating DLL4 in an ERK-dependent manner leading to activation of Notch4 signalling in endothelial cells, although most of the studies focus on latent upregulation of DLL4 in endothelial cells [48] [27]. Due to the importance of DLL4 in viral replication, this redundancy is not surprising. Additionally non-coding RNA expression patterns are cell-type specific, therefore whether circHIPK3 and miR-30c are functionally active in KSHV-infected endothelial cells needs further investigation. Furthermore, exogenous miR-30c was shown to inhibit KSHV replication through targeting of DLL4, although the functional role of endogenous miR-30c has not been previously investigated in KSHV [22].

We demonstrate through flow cytometry analysis the circHIPK3:miR-30c:DLL4 circuit has a role in regulating the host cell cycle during KSHV replication. Although the role of KSHV in the cell cycle during latency is understood, with v-Cyclin playing a key role, the role of the cell cycle during KSHV lytic replication is debated, with lytic replication either requiring entry into S phase or G1 arrest [49] [50] [51]. Our results show that levels of cells in G1 phase slightly increase from 0 to 24 hours while S phase is not significantly altered. This increase of cells into G1 was also noticeable even when treated with G2/M inhibitors. This increase in G1 phase was also accompanied by a loss of cells in G2/M phase with the corresponding peak far less distinct. Other viruses have also been observed promoting G1 phase, notably EBV, a human gammaherpesvirus, has been observed promoting G1 phase during lytic replication [52] [53]. Furthermore, other herpesviruses including HCMV and HSV-1 also require G1 entry for successful lytic replication [54] [55]. Interestingly, these studies found G1 exit into S phase was blocked upon infection, however, several cellular factors found in S phase were still promoted, it is hypothesised these cellular conditions allow successful viral DNA replication using cellular proteins, however, prevent competition from cellular DNA synthesis which occurs in S phase. Further examination is needed to identify whether this occurs during KSHV lytic replication.

Both circHIPK3 and DLL4 depleted cell lines demonstrated the same phenotype using flow cytometry analysis, with a prevention of the G2/M dysregulation seen during lytic replication in scramble control cells, indicative that KSHV-mediated cell cycle dysregulation is being partly blocked. DLL4 has previously been characterised to regulate cyclin and cell cycle factors [27] therefore we hypothesise; that dysregulation of this ncRNA network at any phase prevents KSHV-mediated dysregulation of the cell cycle leading to inhibition of productive viral replication. The regulation of cell cycle through circHIPK3 function aligns with its observed upregulation in many cancers leading to its identification as a pro-oncogenic circRNA, as dysregulation of the cell cycle is a key step in the development of cancer [38]. Interestingly previous research has shown KSHV dysregulates this cell cycle in the development of cancer, however, it has focused on latent dysregulation, and we have therefore shown KSHV dysregulates circHIPK3 during lytic replication to dysregulate the cell cycle [56].

Supporting this hypothesis, we found the dysregulation of cellular cyclins during lytic replication was reduced in circHIPK3 and DLL4 KD cell lines. Additionally in our circHIPK3 and DLL4 KD cells, immunofluorescence confirmed viral replication centres, essential for viral replication form to a much lesser extent resulting in the observed reduction in viral load, once again suggesting that KSHV DNA lytic replication is tied to its dysregulation of the cell cycle. Moreover, to confirm cell cycle dysregulation is important for KSHV replication, lytic cells were treated with various known cell-cycle inhibitors. Arrest in the G2/M phase through RO-3306 and nocodazole treatment, reversing the observed loss of cells in G2/M phase upon typical lytic replication, led to a significant decrease in viral load. In contrast, addition of thymidine, a G1/S exit blocker, had no impact on viral replication, likely due to lytic replication causing a slight increase in cells in G1 phase, implying KSHV-lytic replication requires cells to be in G1 phase. Importantly, addition of thymidine to DLL4 KD cells reversed the anti-viral affect previously observed, due to increases in cells in G1 phase and a decrease in G2/M phase, once again suggesting that DLL4 and circHIPK3 aid viral replication at least partly through cell cycle dysregulation [27].

In summary, elucidating the role of circRNAs within viral infection is still in its infancy, however, in recent years, increasing number of studies have identified viral encoded-circRNAs and also RNA-Seq-based experiments have highlighted aberrant circRNA expression profiles during infection. We have identified the host cell circRNA, circHIPK3, is dysregulated and essential for KSHV lytic replication. Notably, it forms the cornerstone of a novel ncRNA network, regulating the downstream targets of miR-30c and DLL4, which are in turn, all essential for KSHV lytic replication. The finding of this novel network suggests there may be many more roles for circRNAs both in KSHV and in other viruses’ infection.

## Materials and Methods

### Cell culture

TREx-BCBL1-RTA cells, a B cell lymphoma cell line latently infected with KSHV engineered to contain a doxycycline inducible myc-RTA were a gift from Professor JU Jung (University of Southern California). TREx-BCBL1-Rta cells were cultured in RPMI1640 with glutamine (Gibco), supplemented with 10% Foetal Bovine Serum (FBS) (Gibco), 1% P/S (Gibco) and 100 µg/mL hygromycin B (ThermoFisher). HEK-293T cells were purchased from ATCC and cultured in Dulbecco’s modified Eagle’s medium with glutamine (DMEM) (Lonza), supplemented with 10% FBS and 1% P/S. All cell lines tested negative for mycoplasma. Virus lytic replication was induced via addition of 2 µg/mL doxycycline hyclate (Sigma-Aldrich). All cells were cultured at 37 °C at 5% CO_2_. miRNA mimics were transfected at 50 nM using Lipofectamine RNAi Max (ThermoFisher) and left for 24 hours before addition of doxycycline and further experiments. Plasmids were transfected in a 1:2 ratio with Lipofectamine 2000 (ThermoFisher).

### RNA extraction, cDNA synthesis and qPCR

Total RNA was extracted using Monarch Total RNA Miniprep Kit (NEB) as per manufacturer’s protocol. 1 µg RNA was reverse transcribed using LunaScript RT SuperMix Kit (NEB). qPCR was performed using synthesised cDNA, GoTaq qPCR MasterMix (promega) and the appropriate primer. qPCR was performed on Rotorgene Q and analysed by the ΔΔCT method against a housekeeping gene as previously described [57].

For miRNA analysis total RNA was extracted using TRIzol (Invitrogen) as per manufacturer’s protocol. RNA was then DNase treated using DNase I (18068015 Thermo Fisher) as per protocol. 1 µg RNA was reverse transcribed using miScript II RT kit (Qiagen) as per protocol before qPCR using miScript SYBR Green kit (Qiagen) and analysis by the ΔΔCT against SNORD68.

### Plasmids and antibodies

Antibodies used in western blotting are listed below: ORF65 (CRB crb2005224, 1/100), ORF57 (Santa Cruz sc-135747 1/1000), GAPDH (Proteintech 60004-1-Ig 1/5000), GFP (Proteintech 66002-1-ig 1/5000), DLL4 (Proteintech 21584-1-AP 1/200). Ago2 (abcam ab186733) was used in RIPs. RNA pol II (Sigma Aldrich 05-623 1/50) was used in IF.

pVSV.G and psPAX2 were a gift from Dr Edwin Chen (University of Leeds). PLKO.1 TRC cloning vector was purchased from Addgene (gift from David Root; Addgene plasmid #10878) with shRNAs against DLL4 and circHIPK3 cloned into it. psiCheck2 was a gift from Dr James Boyne (Leeds Beckett University). GFP, GFP-ORF50, GFP-ORF57 and GFP-ORF57 RGG1/2 have been described previously [23] [24]. Primers are listed in Table S1.

### Immunoblotting

Cell lysates were separated using 8-12% polyacrylamide gels and transferred to Amersham Nitrocellulose Membranes (GE healthcare) via Trans-blot Turbo Transfer system (Bio-Rad). Membranes were blocked in TBS + 0.1% tween with 5% wt/vol dried skimmed milk powder. Membranes were probed with appropriate primary antibodies and secondary horseradish peroxidase conjugated IgG antibodies at 1/5000 (Dako Agilent). Proteins were detected with ECL Western Blotting Substrate (Promega) or SuperSignal™ West Femto Maximum Sensitivity Substrate (ThermoFisher) before visualisation with G box (Syngene).

### Fluorescence in situ hybridization

TREx-BCBL-1-RTA cells were seeded onto poly-L-lysine (Sigma-Aldrich) coated coverslips and 2 µg/mL doxycycline hyclate (Sigma-Aldrich) was added 3 hours later. FISH was performed 24 hours later using ViewRNA Cell Plus Assay (ThermoFisher) as manufacturer’s protocol. Coverslips were mounted using Vectashield Hardset Mounting Medium with DAPI (Vector laboratories). Images were obtained using a Zeiss LSM7880 Inverted Microscope confocal microscope and processed using ZEN 2009 imaging software (Carl Zeiss) as previously described [58].

### Immunofluorescence

TREx-BCBL-1-RTA cells were seeded onto poly-L-lysine coated coverslips and 2 µg/mL doxycycline hyclate (Sigma-Aldrich) was added 3 hours later. Cells were fixed with for 15 minutes with 4% paraformaldehyde and permeabilised with PBS + 1% Triton. All further incubation steps occurred at 37 °C. Coverslips were blocked for 1 hour with PBS and 1% BSA before 1 hour incubation with the appropriate primary antibody and followed by 1 hour with Alexa-fluor conjugated secondary antibody 488 (Invitrogen 1/500). Coverslips were mounted using Vectashield Hardset Mounting Medium with DAPI (Vector laboratories). Images were obtained using a Zeiss LSM880 Inverted Microscope confocal microscope and processed using ZEN 2009 imaging software (Carl Zeiss) as previously described [58].

### Viral reinfection and viral load assays

TREx-BCBL-1s were induced for 72 hours before being spun down and supernatant was added to 293Ts in a 1:1 ratio with DMEM. 24 hours after addition of supernatant, cells were harvested and RNA extracted, reverse transcribed before qPCR analysis. For viral load TREx-BCBL-1s were induced for 72 hours before harvesting. DNA was extracted using DNeasy Blood and Tissue Kit (Qiagen) as per manufacturer’s instructions before qPCR analysis.

### RNA immunoprecipitations

For biotinylated RIPs HEK-293T cells were seeded and transfected with 20nM biotinylated LNA miR-29b or miR-30c (Qiagen) in combination with 2 μl Lipofectamine RNAi Max (ThermoFisher). 24 hours post transfection, RIPs were performed as per manufacturer’s directions using Dynabeads MyOne Streptavidin T1 (ThermoFisher). RNA was extracted and purified using TRIzol LS (Invitrogen) as per manufacturer’s instructions before analysis via qPCR. Samples were analysed using fold enrichment over % inputs.

Ago2 RIPs were performed on TREx-BCBL-1 RTA cells using EZ-Magna RIP RNA binding Immunoprecipitation Kit (Merck millipore) as per manufacturer’s instructions. RNA was extracted and purified using TRIzol LS (Invitrogen) as per manufacturer’s instructions before analysis via qPCR. Samples were analysed using fold enrichment over % inputs.

For GFP RIPs HEK-293Ts were transfected with 2μg GFP or GFP-ORF57 plasmids with DNA in a 1:2 ratio with Lipofectamine 2000 (Thermo Fisher Scientific). Cells were lysed before incubated with GFP-Trap Agarose beads overnight at 4 °C (Chromotek) and washed as manufacturer’s protocol. Samples were incubated with Proteinase K buffer containing (10 mM Tris pH 7.5, 150 mM NaCl, 0.5 mM EDTA, 10% SDS, proteinase K) for 30 minutes at 55 °C before RNA extracted via TRIzol LS (Invitrogen) as per manufacturer’s instructions before analysis via qPCR. Samples were analysed using fold enrichment over % inputs.

### 3’ UTR Luciferase Assay

The 3’UTR of *DLL4* was identified using NCBI AceView before cloned into psiCheck2 plasmid. HEK-293T cells were transfected with 50 nM miR-30c or scr alongside psi-check2 plasmid. Luciferase reporter assays were performed 24 hours post-transfection using Dual Luciferase Reporter Assay System (Promega) as per the manufacturer’s directions.

### Lentivirus-based shRNA Knockdown

Lentiviruses were generated by transfection of HEK-293T cells seeded in 12-well plates using a three-plasmid system. Per 12-well, 4 µl of lipofectamine 2000 (Thermo Scientific) were used together with 1.2 µg of pLKO.1 plasmid expressing shRNA against the protein of interest, 0.65 µg of pVSV.G, and 0.65 µg psPAX2. pVSV.G and psPAX2 were a gift from Dr. Edwin Chen (University of Westminster, London). Eight hours post-transfection, media was changed with 1.5 mL of DMEM supplemented with 10% (v/v) FCS. Two days post-transfection, viral supernatants were harvested, filtered through a 0.45 µm filter (Merck Millipore) and immediately used for transductions of TREx BCBL1-RTA cells. Cells (500,000) in 12 well plates were infected by spin inoculation for 60 min at 800 x g at room temperature, in the presence of 8 μg/mL of polybrene (Merck Millipore). 3 µg/ml Puromycin (Gibco) was added 48 hours after transduction before KD analysed via qPCR and western blot if appropriate.

### Site directed mutagenesis

Site-directed mutagenesis of GFP-ORF57 for W292A mutant was carried out using QuikChange site-directed mutagenesis kit (Agilent technologies) as per manufacturer’s instructions.

### Cell Cycle Analysis

Cells were seeded, harvested as required, followed by fixing overnight in 70% ethanol at -20 °C. Cells were washed in PBS containing 0.5% BSA, before incubation for 30 minutes at room temperature in PBS containing 0.5% BSA, 5 µg/ml RNase A (ThermoFisher) 25 µg/ml propidium iodide (Sigma). Samples were analysed by CytoFlex S Benchtop Flow Cytometer (Beckman Coulter). For cell cycle drug treatment TREX-BCBL-1 cells were pre-treated for 16 hours with either 10 µM RO-3306, 0.5 µM nocodazole or 2mm thymidine before addition of doxycycline. Viral load was measured after 48 hours.

### miR-Seq Analysis

Total RNA was extracted from TREx-BCBL-1s at 0, 16 and 24 hours post lytic induction. Small RNA libraries were prepared using the TruSeq Small RNA Library Prep Kit (Illumina). Samples were assessed using Agilent High Sensitivity D1000 Screen Tape Station and gel electrophoresis and size based extraction performed. cDNA libraries were analysed via Illumina HiSeq (Illumina) by University of Leeds NGS facility and data deposited at Gene Expression Omnibus (accession number to be assigned).

### RNA-Seq Analysis

Total RNA was extracted from TREx-BCBL-1s at 0 and 18 hours post lytic induction. mRNA sequencing libraries were created using an Illumina TruSeq Stranded mRNA Sample Prep Kit (Illumina). cDNA libraries were generated and analysed using an Agilent Technologies 2100 Bioanalyze. Libraries were sequenced using a HiSeq (Illumina) 75 sequencing platform. Raw reads were processed for RNA expression using standard bioinformatics pipeline. Quality filtered (Q < 20), and adapter trimmed reads (Trimmomatic v0.39) [59] were aligned to the GRCh38/hg38 assembly of the human genome using Bowtie2 (V 2.4.2) or HISAT2 (V 2.1.0) [60] [61] for miRNA and mRNA expressions, respectively. Then the counts in different genomic features were generated using HTSeq (v0.11.1) [62] on human microRNA annotations (miRbase.org) or GRCh38 annotation (GENCODE Release 32). Expression levels were normalised by “counts per million” (CPM). Differential expression (DE) analyses between two KSHV replication times were performed using limma R package. The DE miRNAs and mRNA were defined at adjusted p-value < 0.05. To reduce rate of false discovery rate in the DE analysis we included only transcripts with at least 1 CPM in 3 samples.

### Microarray Analysis

For microarray analysis of miR-30c and miR-29b expression datasets GSE55625 and GSE18437 from NCBI Geo were used and analysed via GEO2R, based on the limma method.

## Acknowledgements

This work was supported in part by grants from the White Rose BBSRC Doctoral Training Partnership in Mechanistic Biology (95519935) MRC DiMeN Doctoral Training Partnership (95505183), BBSRC project grant (BB/T00021X/1) and MRC project grant (MR/V009478/1). We thank Professor Jae Jung, University of Southern California School of Medicine, Los Angeles, for the TREx BCBL1-RTA cells, Dr Edwin Chen, University of Westminster for the lentivirus vectors and Dr James Boyne, Leeds Beckett University for the psiCheck2 plasmid.

