## Supplemental Figures 1-8 for "CircHIPK3 dysregulation of the miR-30c/DLL4 axis is essential for KSHV lytic replication"

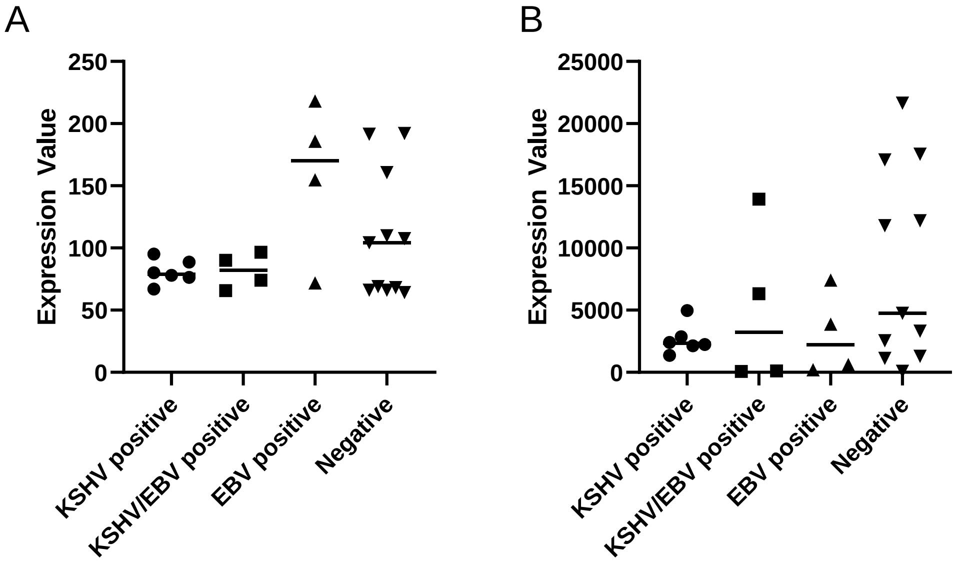


**Supplementary Figure 1: (A)** Scatter plot data of miR-30c from GSE18437 with KSHV positive samples (n=6), KSHV + EBV positive samples (n=4), EBV positive samples (n=4) and KSHV + EBV negative samples (n=11). **(B)** Scatter plot data of miR-29b from GSE18437 with KSHV positive samples (n=6), KSHV + EBV positive samples (n=4), EBV positive samples (n=4) and KSHV + EBV negative samples (n=11).

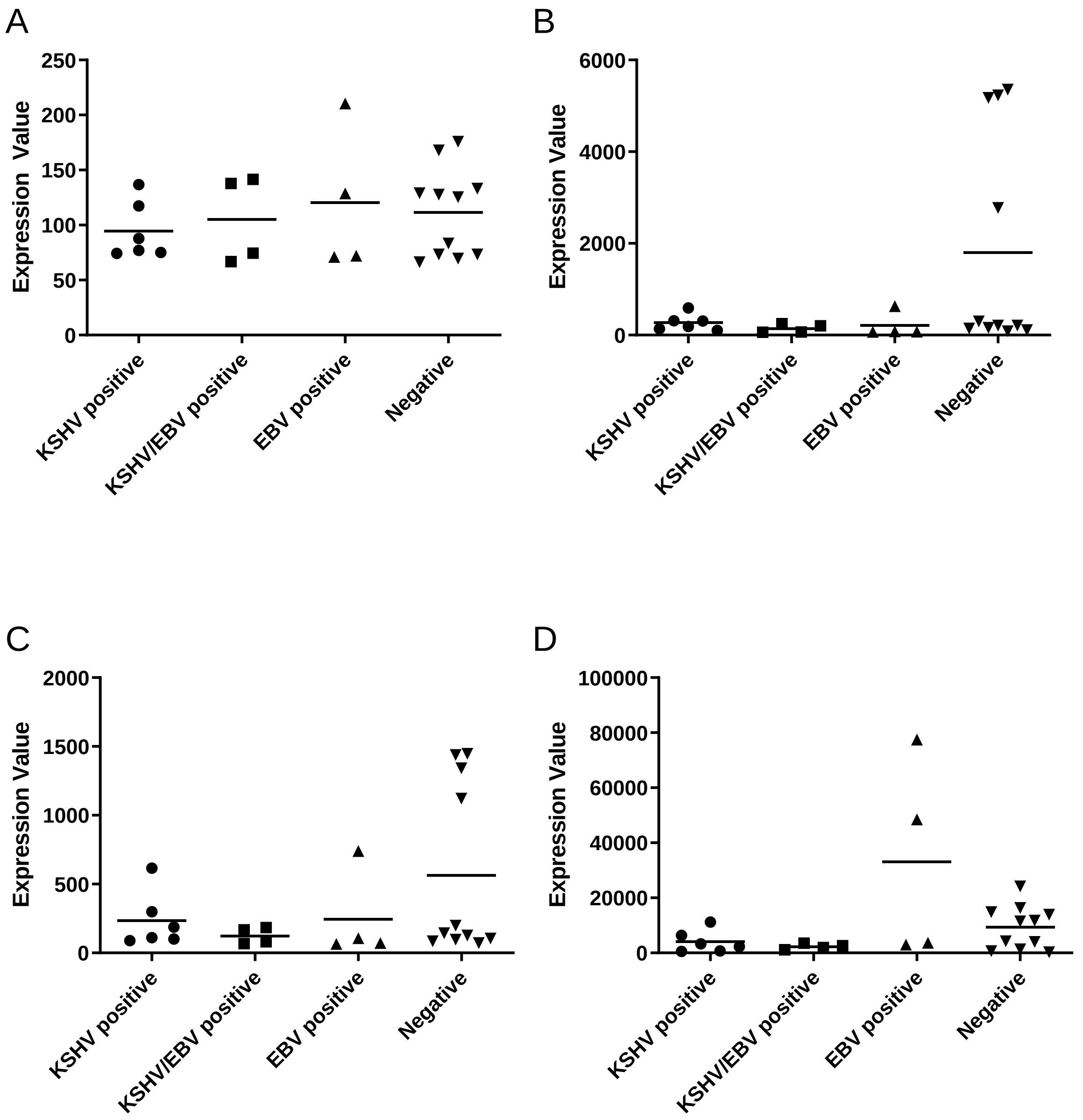


**Supplementary Figure 2: (A)** Scatter plot data of miR-128 from GSE18437 with KSHV positive samples (n=6), KSHV + EBV positive samples (n=4), EBV positive samples (n=4) and KSHV + EBV negative samples (n=11), miR-27 **(B),** miR-23 **(C)** and miR-92 **(D).**

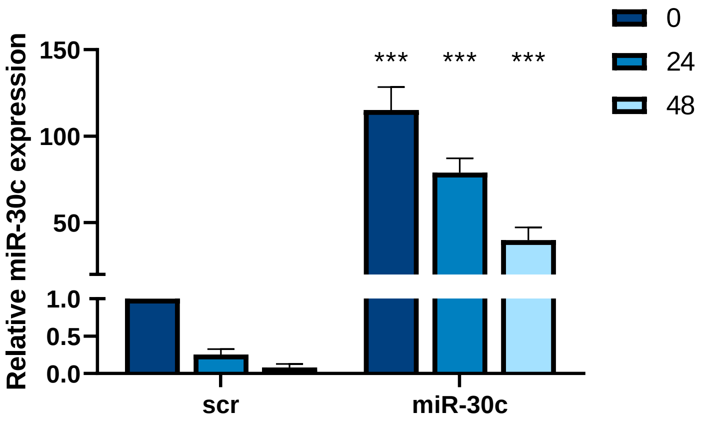


**Supplementary Figure 3:** qPCR analysis of miR-30c levels, cells were transfected with a scrambled control or a miR-30c mimic before lytic replication induced 24 hours after transfection. miR-30c levels were analysed at 0, 24 and 48 hours post lytic induction. SNORD68 was used as a housekeeper (n=3).

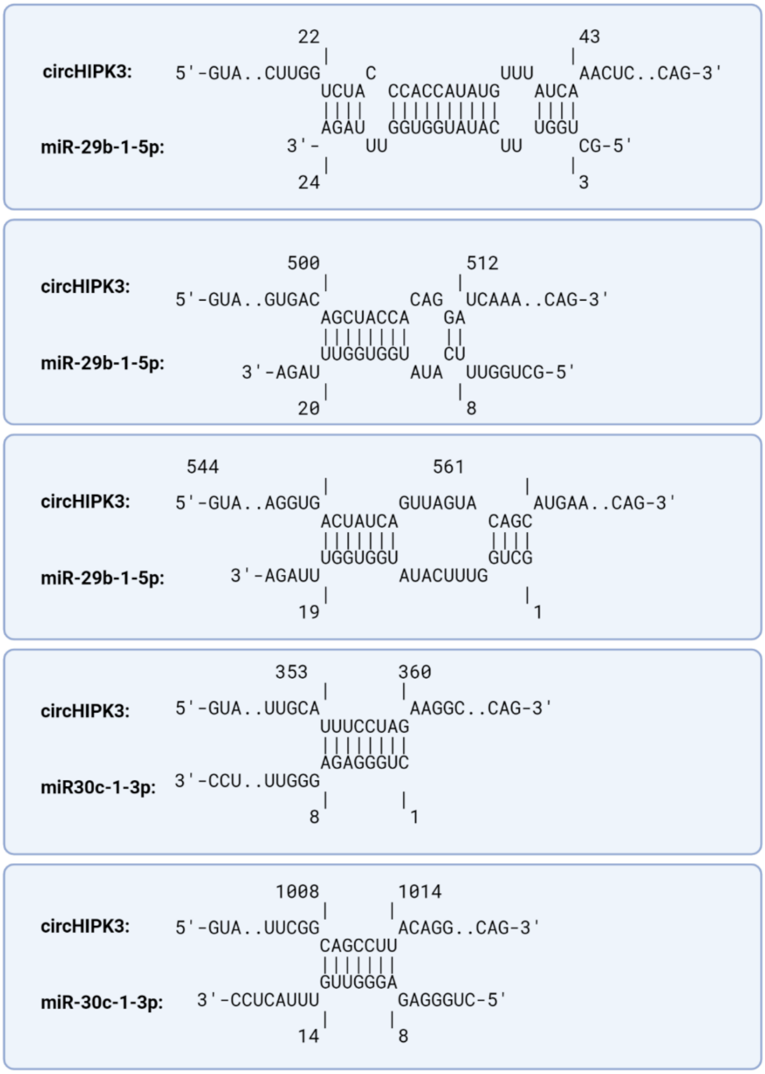


**Supplementary Figure 4:** RNA binding predictions for circHIPK3 and miR-29b/miR-30c

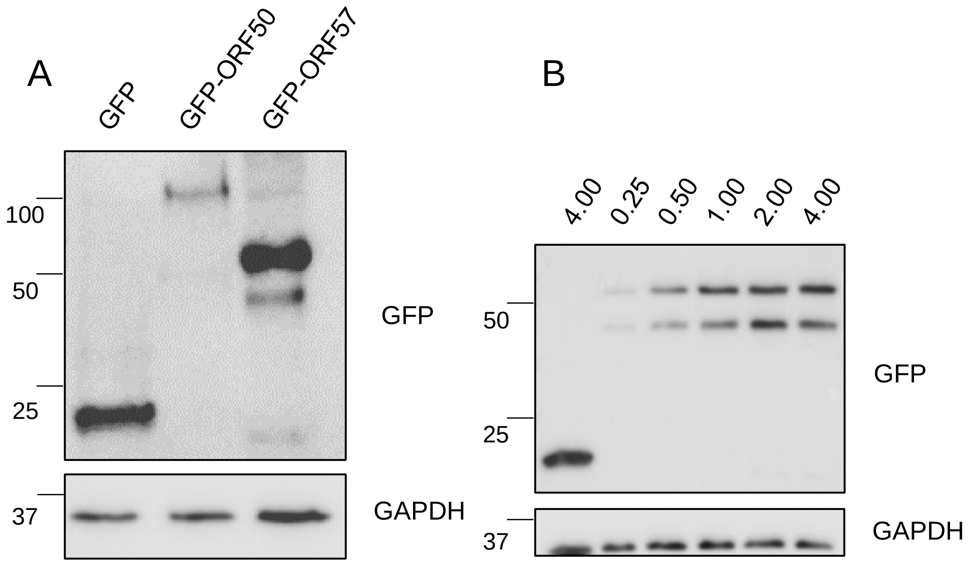


**Supplementary Figure 5: (A)** Western blot for GFP at 24 hours post-transfection in HEK-293Ts of GFP, GFP-ORF50 and GFP-ORF57. GAPDH was used as a loading control. **(B).** Western blot for GFP at 24 hours post-transfection in HEK-293Ts of 4 µg GFP or varying amounts of GFP-ORF57.

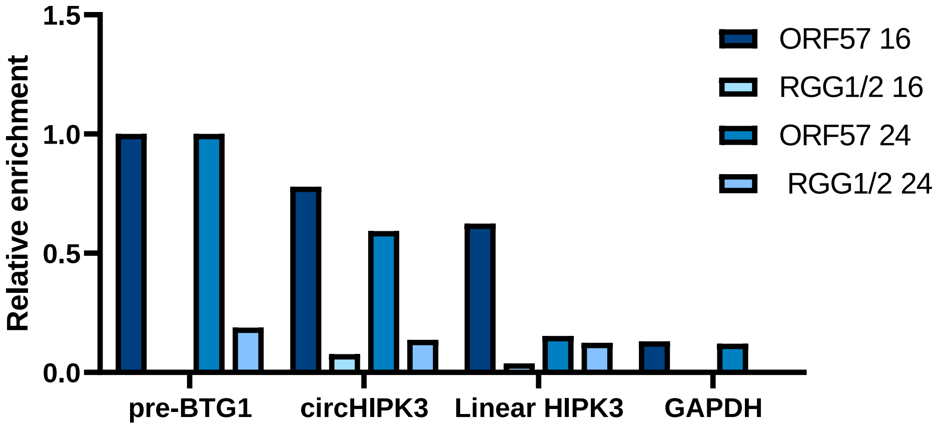


**Supplementary Figure 6:** qPCR analysis GFP RIPs in GFP-ORF57 or GFP-ORF57 RGG1/2 transfected HEK-293Ts at 16 and 24 hours post-transfection respectively n=1 for each time course.

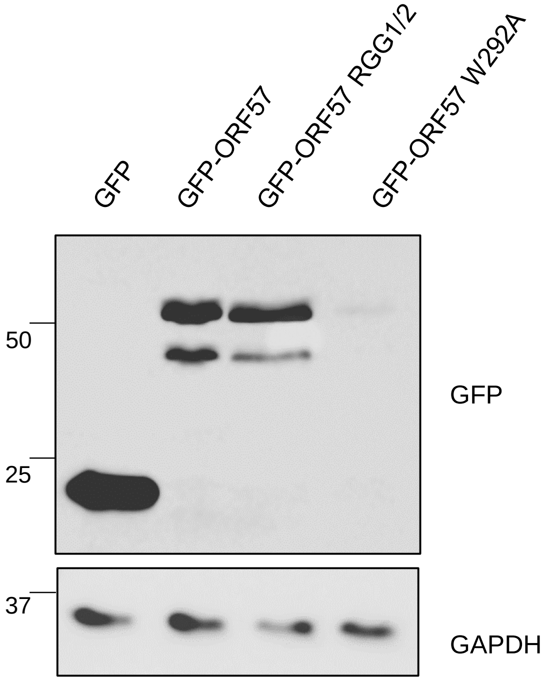


**Supplementary Figure 7:** Western blot for GFP at 24 hours post-transfection in HEK-293Ts of GFP, GFP-ORF57, GFP-ORF57 RGG1/2 and GFP-ORF57 W292A. GAPDH was used as a loading control.

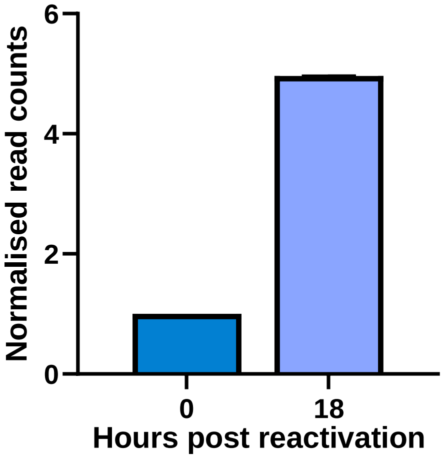


**Supplementary Figure 8:** Levels of DLL4 in mRNA Seq at 0 and 18 hours normalised to read count at 0hrs. N=2

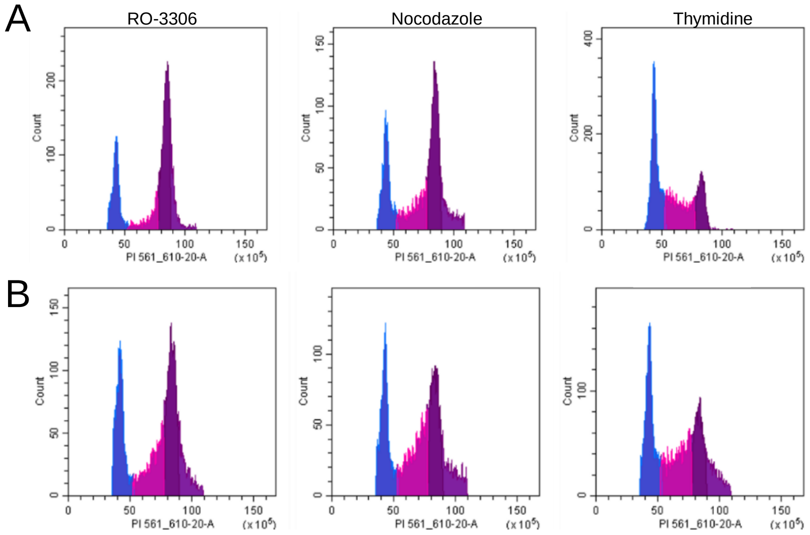


**Supplementary Figure 8:** Representative plots of cell cycle distribution at 0 **(A)** and 24 **(B)** hours post induction in RO-3306, nocodazole and thymidine treated cells, with G1 phase (blue), S phase (pink) and G2/M phase (purple) highlighted.

| Primer Name | Sequence |
|---|---|
| *GAPDH* Forward | TGTCAGTGGTGGACCTGA |
| *GAPDH* Reverse | GTGGTCGTTGAGGGCAATG |
| circHIPK3 Forward | TATGTTGGTGGATCCTGTTCGGCA |
| circHIPK3 Reverse | TGGTGGGTAGACCAAGAGTGGTGA |
| *ORF57* Forward | GCCATAATCAAGCGTACTGG |
| *ORF57* Reverse | GCAGAGAAATATTGCGGTGT |
| *DLL4* Forward | CCCTGGCAATGTACTTGTGAT |
| *DLL4* Reverse | TGGTGGGTGCAGTAGTTG |
| Pri-miR-29b Forward | TGAAAGCAACAGCAGGATGG |
| Pri-miR-29b Reverse | ACCCAAGACAACCTAGAAAGGA |
| Pre-miR-29b Forward | TTGCCACTTGAGTCTGTT |
| Pre-miR-29b Reverse | CTTCTCTACTGTCACCTCTC |
| Pri-miR-30c Forward | CTGATCAACCCTGGACCCTG |
| Pri-miR-30c Reverse | GGTCGCATTCTGTCCGATCT |
| Pre-miR-30c Forward | GTGTAAACATCCTACACTCTCAGC |
| Pre-miR-30c Reverse | TGGCAGAAGGAGTAAACAACCC |
| Linear *HIPK3* Forward | GGCATCAAGAGTGGAATGGA |
| Linear *HIPK3* Reverse | TGATGAATGGTTGGGGATGG |
| Pre-BTG1 Forward | GTCAACGGCACAATTAACAG |
| Pre-BTG1 Reverse | TGCACACAATGGAGTTGATG |
| *CCNB1* Forward | CATGGTGCACTTTCCTCCTT |
| *CCNB1* Reverse | AGGTAATGTTGTAGAGTTGGTGTCC |
| *CCNE1* Forward | CCACACCTGACAAAGAAGATCATCAC |
| *CCNE1* Reverse | GAGCCTCTGGATGGTGCAATA |
| *DLL4* KD 1 siRNA | GCAAGAAGCGCAATGACCACT |
| *DLL4* KD 2 siRNA | GCAGGGAAGCCATGAACAACT |
| circHIPK3 KD siRNA | CTACAGGTATGGCCTCACA |

**Supplementary Table 1:** List of primers and siRNAs
